# Increasing marine trophic web knowledge through DNA analyses of fish stomach contents: a step towards an Ecosystem Based Approach to fisheries research

**DOI:** 10.1101/2023.12.18.572137

**Authors:** Oriol Canals, Anders Lanzén, Iñaki Mendibil, Eneko Bachiller, Xavier Corrales, Eider Andonegi, Unai Cotano, Naiara Rodríguez-Ezpeleta

**Affiliations:** AZTI, Marine Research, Basque Research and Technology Alliance (BRTA). Txatxarramendi ugartea z/g, 48395 Sukarrieta - Bizkaia, Spain; IKERBASQUE – Basque Foundation for Science, Bilbao, Bizkaia, Spain

**Keywords:** blocking primers, DNA metabarcoding, fisheries research, fish diet, stomach content analysis

## Abstract

Multispecies and ecosystem models, which are key for the implementation of ecosystem-based approaches to fisheries management, require extensive data on the trophic interactions between marine organisms, including changes over time. DNA metabarcoding (the simultaneous identification of all taxa in a sample by sequencing a short DNA region) could be used for speeding up large-scaled stomach content data collection. Yet, for DNA metabarcoding to be routinely implemented, technical challenges should be addressed, such as the potentially complicated sampling logistics, the detection of a high proportion of predator DNA, and the inability to provide reliable abundance estimations. Here, we present a DNA metabarcoding assay developed to examine the diet of five commercially important fish, which is feasible to be incorporated into routinary samplings. The method is devised to speed up the analysis process by avoiding the stomach dissection and content extraction steps, while preventing the amplification of predator DNA by using blocking primers. Tested in mock samples and in real stomachs, the method has proven effective and shows great effectiveness discerning diet variations due to predator ecology or prey availability. Additionally, by applying our protocol to mackerel stomachs previously analysed by visual inspection, we showcase how DNA metabarcoding could complement visually based data by detecting overlooked prey by the visual approach. We finally discuss how DNA metabarcoding-based data can contribute to trophic data collection. Our work reinforces the potential of DNA metabarcoding for the study and monitoring of fish trophic interactions and provides a basis for its incorporation into routine monitoring programs, which will be critical for the implementation of ecosystem-based approaches to fisheries management.

## 1. Introduction

The organisms that compose marine ecosystems are highly interconnected and interdependent (Cury et al., 2003), implying that changes in one part of the system have wide-ranging implications elsewhere. In particular, trophic interactions play an important role in marine ecosystem dynamics (Bax, 1998; Frank et al., 2005; Hunsicker et al., 2011) and their understanding is critical in fisheries assessment for ensuring sustainable management (Link, 2002, 2010). Fisheries assessment has traditionally relied on a single-species perspective, i.e., accounting for the dynamics and fishing pressure of a specific species or stock, obviating the ecological (e.g., trophic interactions) and environmental factors (e.g., temperature) affecting population dynamics and the impacts of both manageable and unmanageable pressures (Marshall et al., 2019; Skern-Mauritzen et al., 2016). However, the need of assessing marine resources incorporating such ecosystem processes has become evident and more comprehensive frameworks such as the Ecosystem Approach to Fisheries (FAO, 2003) or the Ecosystem-Based Fisheries Management (EBFM) (Link, 2010) are acknowledged goals in international policy (Juan-Jordá et al., 2018) and at regional/national level (EU, 2013; NOAA, 2016).

Multispecies and ecosystem models contribute to EBFM by allowing to merge available ecological information, considering the relationships between ecosystem compartments and stressors (Christensen & Maclean, 2011; Collie et al., 2016; Townsend et al., 2019). Models can range from including a limited number of species with known, important interactions with the target species (Plagányi et al., 2014), to the inclusion of the entire food web (e.g., Ecopath with Ecosim, Atlantis) (Fulton, 2010). These models usually demand extensive knowledge on trophic relationships, i.e., who eats whom, for which an accurate identification of prey is essential, preferably at species level. Further, some of these models require information about the proportion of each prey in the diet of predators to determine trophic flows (preferably in energetic terms). They also require knowledge about the ontogenetic development stage of both prey and predators (whether the predator and prey are juveniles or adults), and age and/or length of prey. Finally, comprehensive temporal and spatial data of the diet of marine organisms are necessary, as feeding behaviour of marine organisms changes temporally and spatially as a response to different factors, such as changes in prey or predator abundances.

Yet, this data is not easily obtained, which limits the implementation and robustness of multispecies and ecosystem models. To date, data on trophic relationships has been primarily obtained by visually inspecting the prey remains in the stomach contents of predators (Buckland et al., 2017; Hyslop, 1980). However, this approach is highly dependent on the degree of degradation of the prey—only recently ingested organisms or little degraded tissues are normally identified, while rapidly degradable organisms like jellyfish are usually under-detected (Buckland et al., 2017; Pompanon et al., 2012)—and relies on taxonomic expertise and the compromise between the desired taxonomic accuracy and required sample processing time (Nielsen et al., 2018), especially when characterizing the diet composition of planktivorous fish. DNA metabarcoding, i.e., the simultaneous identification of the taxa present in a sample by amplifying a short DNA region (Taberlet et al., 2012), is a promising alternative for complementing visual inspection for stomach contents analysis (Roslin & Majaneva, 2016). This approach allows for a more accurate and broader detection and identification of prey diversity compared to visual assessments, especially for preys in highly digested state and particularly for organisms that degrade rapidly, such as gelatinous organisms (Albaina et al., 2016; Bachiller et al., 2021; Bessey et al., 2019; Günther et al., 2021; Maes et al., 2022; Riccioni et al., 2018). In addition, it does not require extensive taxonomic expertise and is more prone to be standardised and automated than visual inspection (Traugott et al., 2021). This method is more cost-effective than visual approaches, and especially suitable to be incorporated in routine data collection programs aiming to obtain comprehensive temporal and spatial data (Thomsen & Willerslev, 2015).

In the context of stomach content analysis, a main shortcoming of DNA metabarcoding is related to the inability of the method to avoid sequencing of predator DNA, which is more abundant and intact than prey DNA (Pompanon et al., 2012; Vestheim & Jarman, 2008). This leads to a great majority of the sequences obtained (also named reads) belonging to the predator instead of prey, being thus uninformative and decreasing the efficiency of the approach (Bessey et al., 2019; Piñol et al., 2014). This can be overcome by using ‘blocking primers’, which impede the amplification of the predator DNA (Vestheim & Jarman, 2008). Blocking primers have been already used in fish diet studies (Bachiller et al., 2020; Devloo-Delva et al., 2019; Su et al., 2018); however, it should be considered that it may come with the cost of undesired co-blocking in the amplification of prey species (Piñol et al., 2015).

In European marine regions, DNA metabarcoding has only been applied to assess the diet of few commercially relevant fish species: European hake (*Merluccius merluccius*) (Gül et al., 2023; Riccioni et al., 2018), polar cod (*Boreogadus saida*) (Maes et al., 2022), Atlantic bluefin tuna (*Thunnus thynnus*) (Günther et al., 2021), European sardine (*Sardina pilchardus*), European anchovy (*Engraulis encrasicolus*), Round sardinella (*Sardinella aurita*), and European sprat (*Sprattus sprattus*) (Albaina et al., 2016; Bachiller et al., 2020; Bachiller et al., 2021), from which only two studies have been conducted in ICES ecoregions (Albaina et al., 2016; Maes et al., 2022). Although DNA metabarcoding has great potential for providing large-scaled fish trophic data, its incorporation into data collection routinary programs is hampered by, among other reasons, the lack of standardised method.

Here, we aim at facilitating diet data collection programs by developing and testing a standardised DNA metabarcoding protocol for prey identification that avoids the need of dissecting and extracting the stomach contents. The protocol has been developed for five highly commercial fish species in the Bay of Biscay, and its efficiency tested using both mock stomach samples and real stomachs of the above-mentioned predators. Also, we have compared the performance of both visual inspection and DNA metabarcoding approaches for assessing the diet of Atlantic mackerel in the Bay of Biscay. The presented protocol represents a significant advance towards the high-throughput and routine analysis of stomach contents to support the ecosystem-based fisheries management.

## 2. Material and methods

### 2.1. Ethical Statement

Samples were collected during annual multidisciplinary oceanographic surveys carried out within the EU Data Collection Framework (DCF) following standard methodologies commonly used in scientific monitoring. Marine species were not exposed to any unnecessary pain, injuries, or suffering. Also, special attention was given to the avoidance of accidental catches of sensitive or endangered species (marine mammals, seabirds, etc.).

### 2.2. Sample collection and preservation

Tissue samples from marine organisms naturally found in the Bay of Biscay, including 10 bony fish (Actinopterygii), 3 cartilaginous fish (Elasmobranchii), 2 polychaetes (Annelida), and one decapod (Crustacea), bivalve (Mollusca) and cephalopod (Mollusca) species were obtained from the AZTI tissue bank (see Table S1).

Adult specimens of European hake, European anchovy, European sardine, and Atlantic horse mackerel, and adult, juvenile and larvae individuals of Atlantic mackerel were obtained from different scientific surveys operating in the Bay of Biscay (fish size, survey, date, and site of capture is provided in Tables S2). On-board, the stomach of each adult or juvenile specimen was extracted, and the excised stomachs and the whole larvae were stored in 96% ethanol at −20 °C.

#### Stomach content characterization under the microscope

The stomachs of the Atlantic mackerel individuals used for the comparison between visual and molecular approaches (41 adults and three juveniles) were thawed and dissected at the laboratory. Gut contents were analysed individually, with no subsampling, under a NIKON SMZ1270 stereomicroscope with 20–80 × amplification. The microscope analysis was conducted in a ‘clean room’ and with an air extractor placed 20–30 cm above the petri plate with the gut samples, following the standard procedures to avoid contamination during sample processing; glassware, bench, microscope slide and dissection equipment (stainless-steel scissors, scalpel, and lancet) were rinsed with 96% ethanol prior to each gut content analysis (Cole et al., 2014). To exclude bias caused by different rates of digestion and cod-end feeding (Hyslop, 1980) only material contained in the stomachs was considered, whereas the contents of the intestine and oesophagus were discarded. During processing, stomach contents were carefully taken apart and all identifiable prey counted and specified to the lowest possible taxonomic group, not including broken parts of appendixes when quantifying. After the microscope analysis, the stomach contents were collected again and stored in 96% ethanol.

### 2.3. DNA extraction

Both the entire stomachs (6 European anchovy and Atlantic horse mackerel, 8 European hake, 9 European sardine, and 14 Atlantic mackerel) and the 44 previously extracted and visually inspected Atlantic mackerel stomach contents were separately homogenized in a blender for few minutes. Between each stomach, the blender was cleaned with 10% bleach, distilled water and 70% ethanol to avoid cross-contamination. All samples (also including the Atlantic mackerel larvae) were centrifuged at 3,000 g for 3 min to remove the ethanol before the DNA extraction step. Genomic DNA was extracted from 20 mg of the homogenised stomachs using the QIAamp DNA Mini Kit (QIAGEN) following manufactureŕs instructions for “DNA Purification from Tissues”, incubating with Proteinase K for overnight instead of 1-3h. DNA from mock stomachs and from larvae was extracted from 20 mg of mock sample and from the whole larvae, respectively, using the Wizard® Genomic DNA Purification kit (Promega, WI, USA) following manufacturer’s instructions for “Animal Tissue”. Extracted DNA was suspended in Milli-Q water and its concentration was determined with the Quant-iT dsDNA HS assay kit using a Qubit® 2.0 Fluorometer (Life Technologies). DNA integrity was assessed by electrophoresis, migrating about 1-2µl of template DNA using GelRed™-stained DNA on an agarose 1.0% gel.

### 2.4. Design of blocking primers

Blocking primers (BP) are short, species-specific sequences with a modification at the 3’ end that specifically prevents the DNA amplification of the target species (Vestheim & Jarman, 2008) by impeding the extension of the polymerase and thus the generation of double-stranded amplicons. Here, a 313 bp fragment of the mitochondrial cytochrome c oxidase I (COI) gene was targeted using the degenerated metazoan universal primers mlCOIintF (Leray et al., 2013) and dgHCO2198 (Meyer, 2003) on which blocking primers were designed as follows. First, COI sequences of each of the species under study were retrieved from GenBank and aligned to generate a consensus sequence per specie using Bioedit (Hall, 1999). Then, amplification primer (AP) sequences were located on the consensus sequence and a specific BP was designed for five species in the Bay of Biscay—European hake, European anchovy, European sardine, Atlantic mackerel and Atlantic horse mackerel—so that it i) overlaps with at least 10 bases with the forward or reverse amplification primers, ii) maximizes the number of matches to the species of interest and iii) minimizes the number of matches to other species. The generated BP are shown in Table 1.

**Table 1.**
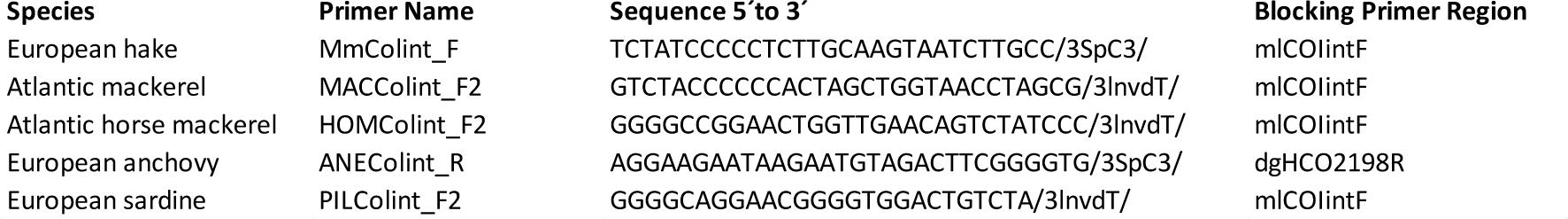
Blocking primers generated for European hake, Atlantic mackerel, Atlantic horse mackerel, European anchovy, and European sardine.

### 2.5. Mock stomach samples construction

Mock samples were constructed by mixing equal proportions of DNA of each species in Table S1, and the mixtures were then combined with each predator DNA at different percentages (100%, 70%, 30%, 0%) and divided into two subsamples to create technical replicates. Four different AP:BP (amplification to blocking primer) ratios (no BP, 1:2, 1:5, and 1:10) were then used to amplify each mock stomach sample (see PCR amplification details below), obtaining a total of 32 different samples for each target predator species (see Table S3).

### 2.6. PCR amplification, library preparation, and sequencing

PCR amplification of extracted DNA was carried out using the mlCOIintF and dgHCO2198R primer pair targeting a 313-bp long, highly variable region of the COI gene. Each sample was amplified three times in a total volume of 20 µl using: i) 10 µl of Phusion® High-Fidelity PCR Master Mix (Thermo Scientific), ii) 0.4 and 0.5 µl of each annealing primer (10 µM) in mock and real stomach samples, respectively, iii) 0, 1, 2.5 or 5 µl of blocking primer (10 µM) in mock samples depending on the AP:BP ratio used (Table S3) and 0.8, 1, 1.2, 2 or 4 µl of blocking primer (10 µM) for obtaining different AP:BP ratios, selected based on PCR visualization on agarose gels, for real stomach samples (Table S2), and iv) 4 µl of genomic DNA (5 ng/µl), with the following PCR conditions: an initial 3 min denaturation step at 98 °C; 27 cycles of 15 sec at 98 °C, 30 sec at 46 °C and 45 sec at 72 °C; and a final 5 min extension at 72 °C. The three replicates were pooled, purified using AMPure XP beads (Beckman Coulter), quantified using the Quant-iT dsDNA HS assay kit using a Qubit® 2.0 Fluorometer (Life Technologies) and used as template for the generation of dual-indexed amplicons in a second PCR round following the “16S Metagenomic Sequencing Library Preparation” protocol (Illumina) using the Nextera XT Index Kit (Illumina). Multiplexed PCR products were purified again using the AMPure XP beads, quantified using Quant-iT dsDNA HS assay kit and a Qubit 2.0 Fluorometer (Life Technologies), normalized to equal concentration and sequenced using the 2 x 300 paired-end MiSeq (Illumina).

### 2.7. Raw read processing, statistical analysis, and data visualization

Initial processing of raw sequence data was performed as described in Hestetun et al. (2021). Briefly, read pairs were merged using *vsearch* v2.7.1 (Rognes et al., 2016) allowing for up to 40 mismatches. Using *cutadapt* v1.15 (Martin, 2011), all merged sequences that lacked complete forward and reverse primers were discarded allowing for up to two mismatches, and the primers were trimmed. Remaining amplicon sequences longer than 333 bp, shorter than 274 bp or with one or more incorrect base calls according to *vsearch* were discarded. SWARM v3.0.0 (Mahé et al., 2014) was used to group dereplicated remaining high-quality sequences into OTUs using default settings (-d1, fastidious clustering and breaking initial OTUs likely to represent different sequence variants), followed by removal of singletons, reference based and *de novo* chimera filtering, using *vsearch*. Dereplication and conversion of SWARM output to an OTU contingency table was done using scripts adopted from SLIM (Dufresne et al., 2019). Finally, LULU (Frøslev et al., 2017) was used for post-clustering correction, with a 97% similarity cut-off, followed by taxonomic assignments of OTUs using CREST v3.2.1 (Lanzén et al. (2012); https://github.com/lanzen/CREST) using BOLD as reference database (Ratnasingham and Hebert (2007); accessed February 2018 and adapted to CREST). OTUs and reads were collapsed by taxonomy into phylotypes, and only those belonging to Metazoa and successfully classified to Phylum rank (representing 99.1% of reads) were kept in further steps. Phylotypes classified as Insecta, Mammalia, Aves, or Arachnida were excluded (42 OTUs representing 0.2% of total reads). Also, five phylotypes (representing 13.6% of reads) belonged to Nematoda, comprising parasitic organisms in the Bay of Biscay such as *Anisakis* spp., which were not considered as preys and were therefore excluded from further analysis. Samples with less than 5,000 reads after filtering were removed (leaving 268 out of 283 samples). Finally, because of minimum differences between the contents of those samples corresponding to the same stomach but analysed using a different AP:BP ratio, they were merged into one unique integrated sample by summing up the absolute number of reads.

All statistical analysis and plots were carried out within the R statistical environment (R version 3.6.2; R Core Development Team (2013)). Co-blocked species in mock samples due to the use of BP were identified based on the standardised residuals from chi square tests (*chisq.posthoc.test* function, chisq.posthoc.test package version 0.1.2; Ebbert (2019)). Ordination of stomach contents was carried out using nonmetric multidimensional scaling analysis (NMDS; *metaMDS* function, vegan package version 2.5-4; Oksanen et al. (2019)) based on Bray-Curtis dissimilarity matrices (*vegdist* function, vegan package). ANOSIM (analysis of similarity; Clarke (1993)) was used to test whether there were significant differences between predefined sample groups (*anosim* function, vegan package). Samples were mapped using the ggmap package (version 3.0.0).

## 3. Results

### 3.1. Blocking primer performance

The effectiveness of the generated BP was tested on mock stomach samples under different AP:BP ratios (no BP, 1:2, 1:5, 1:10) and proportions of prey DNA (0%, 30%, 70%, 100%) (Figure 1A). All BP succeeded in blocking predator DNA amplification when prey DNA was included, regardless of the proportion of prey DNA and the AP:BP ratio (as expected, predator DNA was normally amplified when no BP was used). In mock samples without prey DNA (i.e., simulating empty stomachs), BP for Atlantic mackerel, Atlantic horse mackerel, and European hake were able to hamper predator DNA, especially at low AP:BP ratios (1:5 and 1:10). BP for European anchovy and European sardine were unable to block predator DNA amplification when it was the only source of DNA. Note that some mock samples without prey DNA presented DNA from other species than the predator (probably due to contamination while constructing the mock samples), the amplification of which increased at low AP:BP ratios because this DNA represented a more available source of DNA for the primer binding than predator DNA. Also, although presenting equal proportions of DNA, the species composing the mock construction were differently amplified in the PCR amplification, which indicates existence of PCR bias due to primer efficiency.

**Figure 1.**
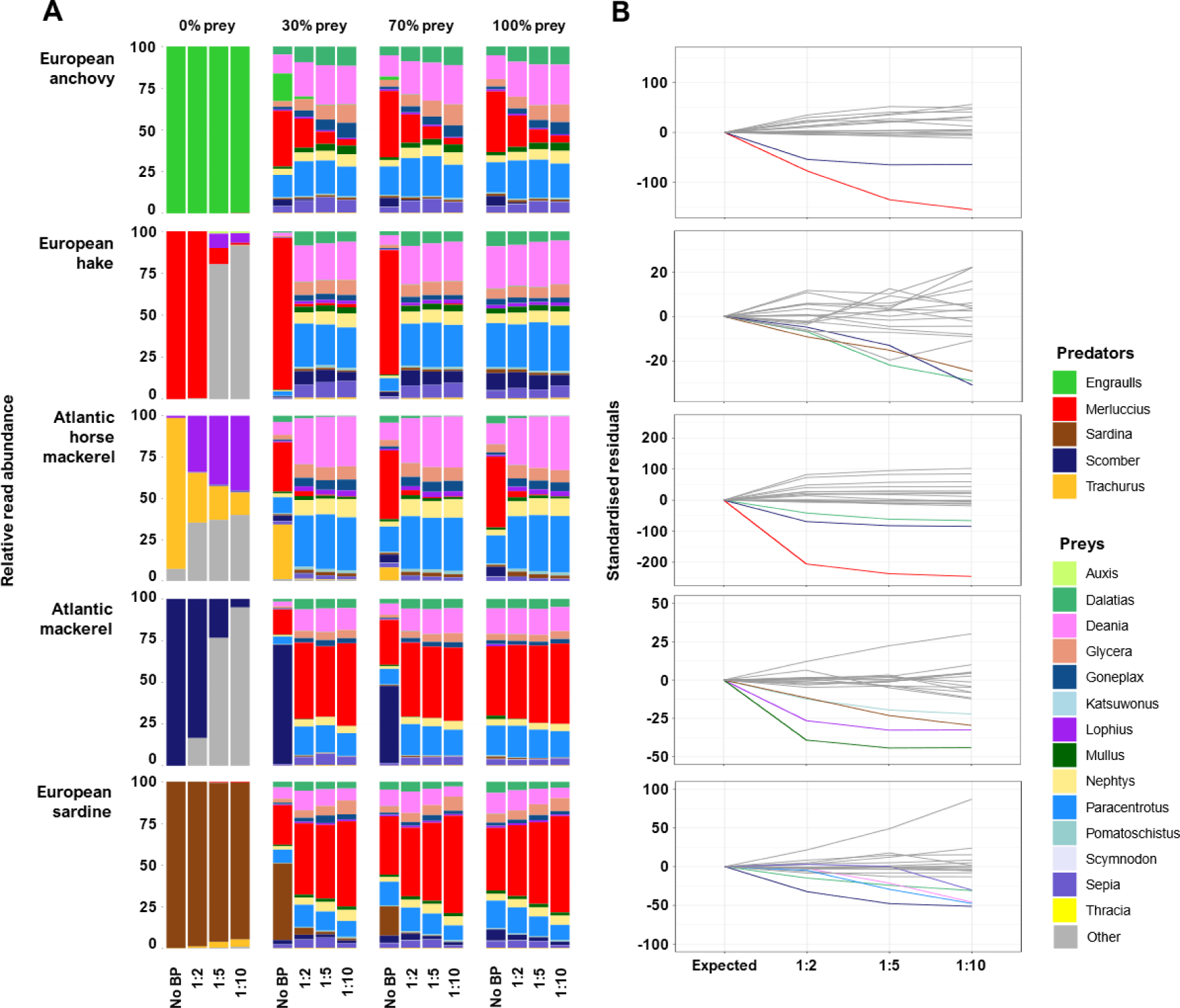
**A.** Relative read abundance of the different species conforming the mock stomach samples for European anchovy, European hake, Atlantic horse mackerel, Atlantic mackerel, and European sardine. **B.** Standardised residuals of chi square test between No BP and AP:BP ratios of 1:2, 1:5, and 1:10 for the 100% prey mock samples (only the species with the lowest standardised residuals, indicating co-blocking, are coloured).

Co-blocking effects by the BP on the species composing the mock samples were identified based on the standardised residuals of the chi square test (Figure 1B). Strong co-blocking of European hake and Atlantic mackerel was observed using the BP designed for European anchovy and Atlantic horse mackerel, as well as co-blocking of *Dalatias licha* by Atlantic horse mackerel BP. BP designed for European hake moderately co-blocked European sardine, Atlantic mackerel, and *Dalatias licha*. Atlantic mackerel BP prevented the amplification of *Lophius piscatorius*, *Mullus surmuletus*, and to a lesser extent *Katsuwonus pelamis* and European sardine. Finally, Atlantic mackerel, *Paracentrotus lividus*, *Deania calcea*, *Dalatias licha*, and *Sepia officinalis* were also somewhat co-blocked using the BP designed for European sardine. The number of mismatches between the designed BP and the sequences of the co-blocked species ranged from a minimum of 2 (for the elasmobranch *Dalatias licha* with the BP designed for Atlantic horse mackerel) to a maximum of 9 (*Paracentrotus lividus* with the BP designed for European sardine) (Figure S1). Notably, most co-blocked species were still detected even at the lowest AP:BP ratio.

### 3.2. DNA metabarcoding analysis of real stomach samples

Our protocol was applied to real stomachs (and few larvae) of five highly commercial species of the Bay of Biscay: European anchovy, European sardine, Atlantic horse mackerel, European hake, and Atlantic mackerel (stomachs and larvae). A total of 11,180,655 reads were obtained after the raw read processing corresponding to 370 different phylotypes, most of which (82.4%, representing 77.5% of reads) were successfully classified at the genus or the species level. Even with the use of BP we still obtained reads belonging to the predator species, although less than 25% in most cases, with the only exception of European sardine, where 64.0±26.5% of reads belonged to the predator (Figure 2A). The number of preys detected within the stomachs of the different predators varied across both species and individuals and suggested a restricted prey spectrum for the demersal European hake compared to the pelagic species (Figure 2B).

**Figure 2.**
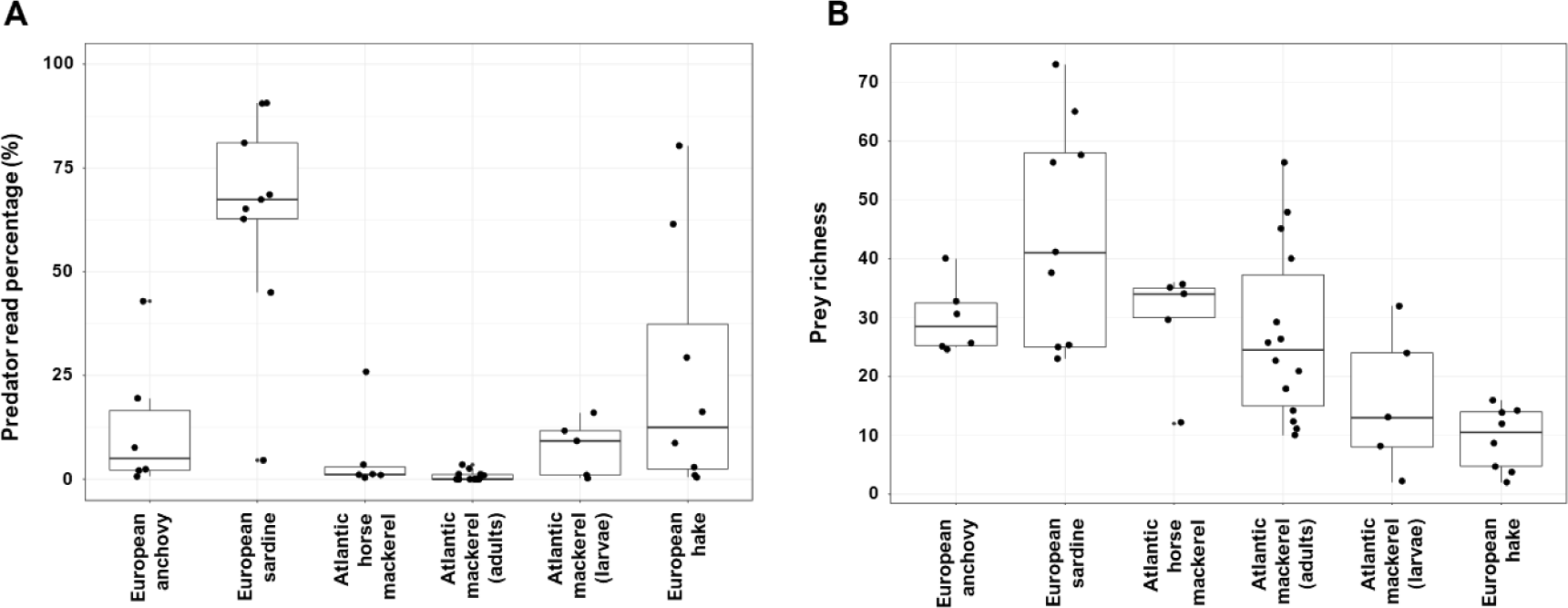
Percentage of predator reads (**A**) and number of prey (**B**; read number rarefied to 3,060 reads) detected in the stomach contents of European anchovy (N=6), European sardine (N=9), Atlantic horse mackerel (N=8 and 7 in A and B, respectively*), Atlantic mackerel (adults and larvae; N=14 and N=5, respectively), and European hake (N=8) in stomachs collected across the Bay of Biscay. The line in the boxes indicates the median. * In **B**, one Atlantic horse mackerel stomach was excluded due to having less than 3,060 reads.

Our results also showed differences in the diet composition among predators (Figure 3A). Arthropoda appeared as the main food source for European anchovy, European sardine, Atlantic horse mackerel, and Atlantic mackerel larvae, and as a prominent prey for Atlantic mackerel adults; but it represented a small proportion of the diet of European hake individuals. Instead, European hake diet was mainly composed by vertebrate preys (mostly Actinopterygii), and to a lesser extent by Echinodermata, Mollusca, and the above-mentioned Arthropoda. Vertebrates also appeared as a relevant input in the diet of Atlantic mackerel but were little abundant in the stomachs of Atlantic horse mackerel, European sardine, and European anchovy. Gelatinous organisms (i.e., cnidarians and tunicates) were detected in the stomachs of all species and represented the second main food source for Atlantic mackerel and European sardine. Differences in the composition of stomach contents were not only observed between but also within each predator species, i.e., different individuals from the same species showed different stomach content composition (Figure 3B). A high proportion of Vertebrata reads (21.5%) was observed in the diet of Atlantic mackerel larvae due to one unique larvae specimen presenting a high proportion of European sardine-belonging reads (Figure 3B). Excluding this specimen, Arthropoda represented 96.9% of Atlantic mackerel larvae diet.

**Figure 3.**
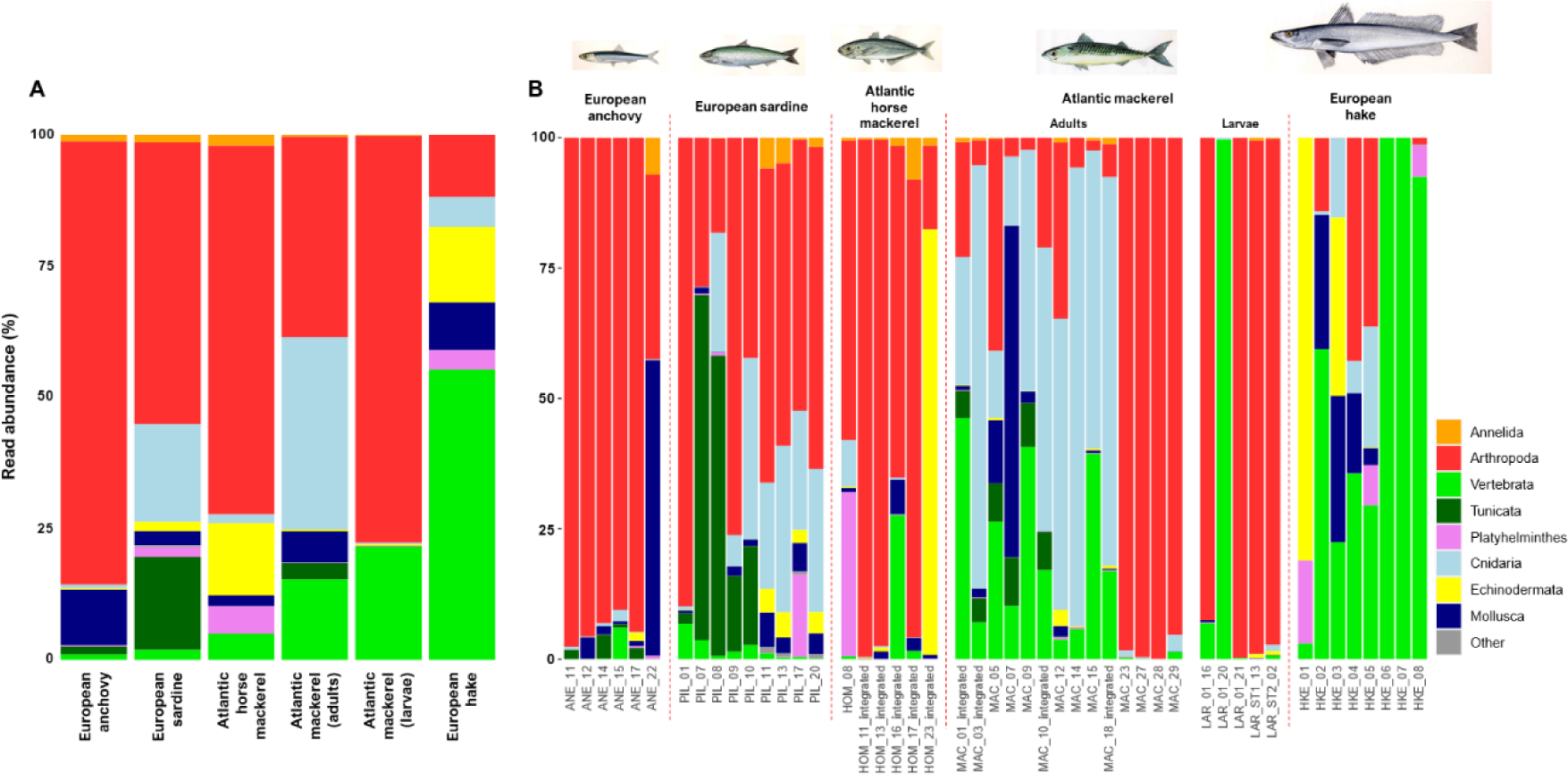
Composition of the stomach contents of European anchovy, European sardine, Atlantic horse mackerel, Atlantic mackerel (adults and larvae), and European hake in the Bay of Biscay. **A:** Overall diet composition for each species; **B:** diet composition for each specimen. Colours in the bars indicate type of prey grouped by Phylum excepting Chordata preys, which have been coloured at the Subphylum level (Vertebrata and Tunicata).

Dietary differences between species were also reflected by an NMDS analysis (Figure 4A). The ordination of stomach content samples of adult specimens clearly differentiated the diet of European hake, the only demersal fish, from pelagic species (i.e., European anchovy, European sardine, Atlantic horse mackerel and Atlantic mackerel) (ANOSIM R statistic: 0.887, p-value <0.001). The composition of stomach contents of pelagic species was not determined by the species (ANOSIM R statistic: 0.096, p-value: 0.081) but by the haul, i.e., the site they were captured at (ANOSIM R statistic: 0.386, p-value <0.001) (Figure 4B).

**Figure 4.**
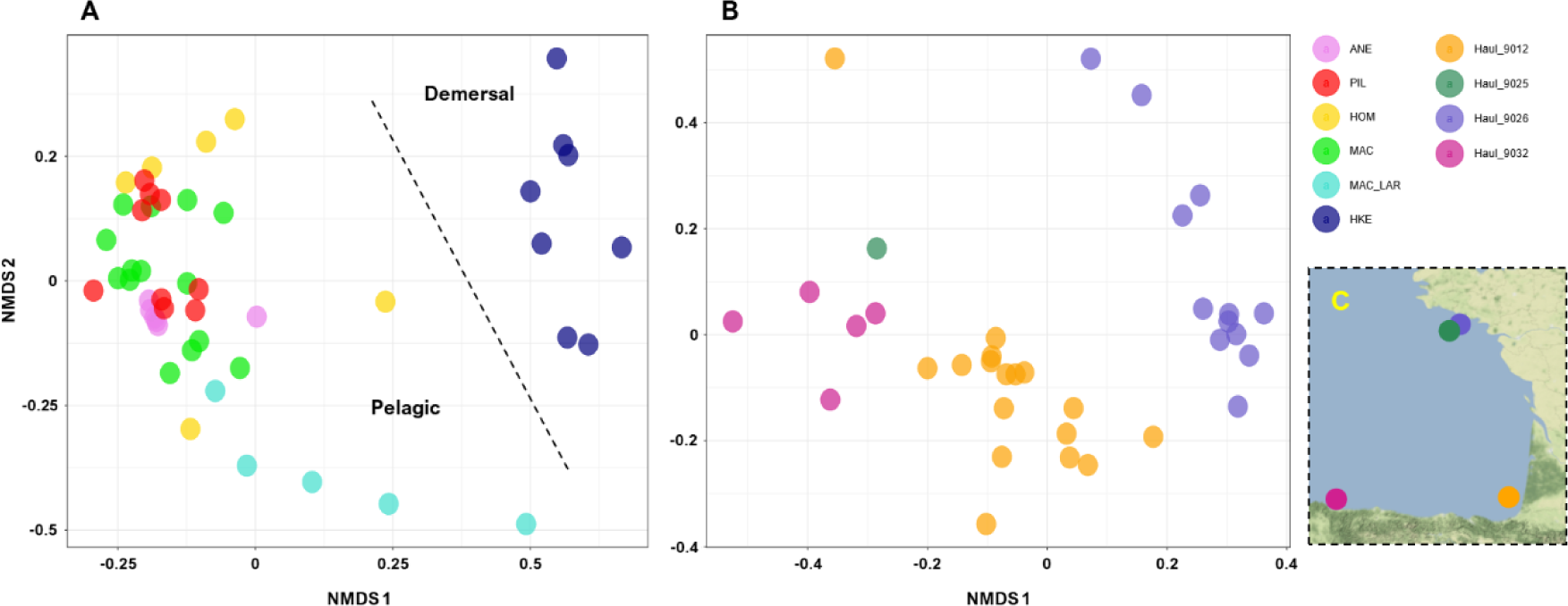
**A:** Ordination of stomach contents by NMDS analysis based on Bray Curtis distances including both pelagic (European anchovy, European sardine, Atlantic horse mackerel, Atlantic mackerel) and demersal (European hake) species, and larvae specimens from Atlantic mackerel. Colour indicates the different species. ANE: European anchovy, PIL: European sardine, HOM: Atlantic horse mackerel, MAC: adult Atlantic mackerel, MAC_LAR: Atlantic mackerel larvae, HKE: European hake. Stress: 0.173. **B:** Ordination of stomach contents only for pelagic fish (i.e., excluding European hake and Atlantic mackerel larvae). Colour indicates the haul from which specimens were collected. Stress: 0.161. **C:** Map of the Bay of Biscay showing the stations of the hauls.

### 3.3. Comparison of visually and DNA metabarcoding inferred Atlantic mackerel diet

The stomach contents of 44 Atlantic mackerel individuals were analysed through visual inspection and DNA metabarcoding. 4,484,508 reads were obtained corresponding to 281 taxonomic categories, 221 of which were successfully classified at the genus or species level (77.3% of reads). On the other hand, visual inspection provided 21 different prey categories, from which 6 (28.6%) could be identified to the species or genus level. Remarkably, all these six visually detected species and genera were also retrieved by DNA metabarcoding. Both methods agreed in pointing to Arthropoda and Vertebrata as the main food source of Atlantic mackerel in the Bay of Biscay (Figure 5A). Yet, noticeable differences were observed regarding the detection and quantification of the other taxonomic groups. The most obvious example is cnidarians, which accounted for an average of 12.3% of DNA metabarcoding reads and were not detected in the stomach remains by visual inspection. Also, molluscs, annelids, and echinoderms were almost only detected by the DNA metabarcoding approach. Finally, ordination of samples by NMDS analysis showed that the stomach contents of Atlantic mackerel individuals were clearly grouped by haul location in both DNA metabarcoding (Figure 5B; ANOSIM statistic R: 0.8, p-value: 0.0001) and visual inspection (Figure 5C; ANOSIM statistic R: 0.606, p-value: 0.0001) approaches.

**Figure 5.**
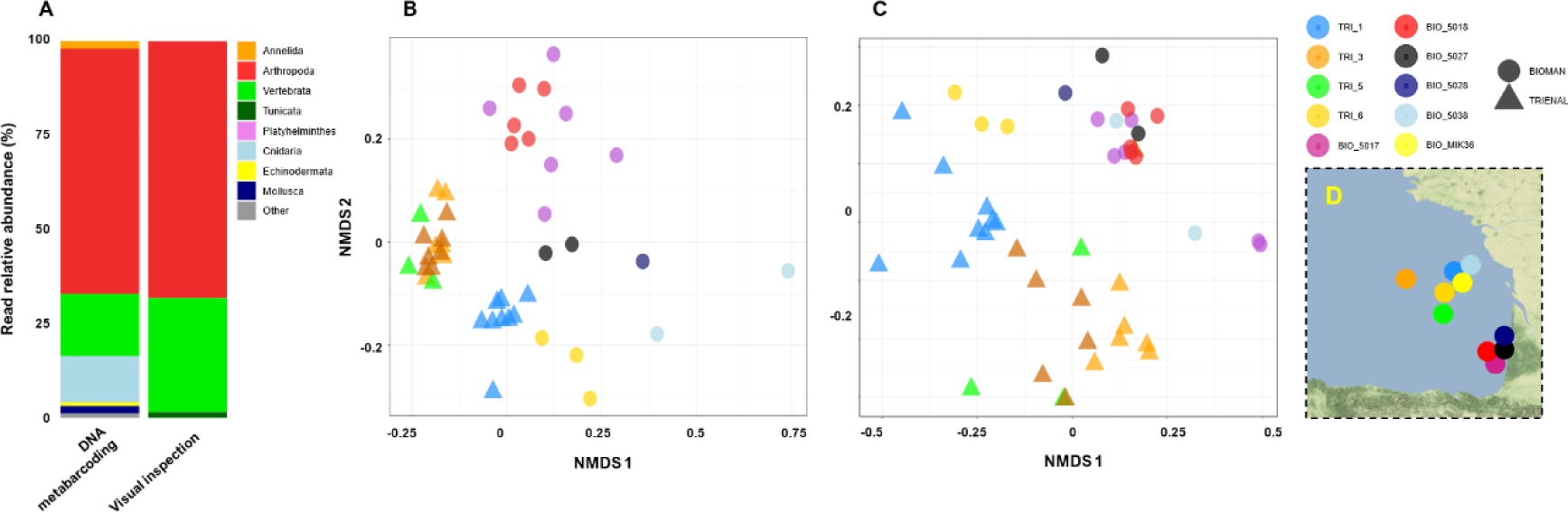
**A:** Composition of the stomach contents of Atlantic mackerel specimens based on DNA metabarcoding and visual inspection (N=44). **B-C:** Ordination of Atlantic mackerel stomach contents by NMDS analysis based on Bray Curtis distances for DNA metabarcoding (**B**; stress: 0.156) and visual inspection-based data (**C**; stress: 0.137). Color indicates the haul and shape indicates the survey from which specimens were collected. **D:** Map of the Bay of Biscay showing the stations of the hauls.

## 4. Discussion

Application of EBFM is still scarce (Marshall et al., 2019; Skern-Mauritzen et al., 2016), partly hampered by the lack of precise and temporally and spatially comprehensive trophic data required by the multispecies and ecosystem models. In the European Union, stomach collection by member states has been recently supported by the EU MAP Data Collection Framework (EU, 2021). When analysed, stomach contents are typically inspected visually. Here, we propose an alternative, complementary method based on DNA metabarcoding to speed up obtaining trophic data of small to medium pelagic and demersal commercial fish, which is suitable to be incorporated into routinary samplings. The main novelty of the method is the processing of the entire stomachs instead of dissecting them for extracting the contents, which, together with the use of blocking primers (Vestheim & Jarman, 2008), is expected to reduce the time needed for the stomach processing, limit cross-contamination between samples, and maximise amplification of prey DNA over predator DNA.

### 4.1. Use of blocking primers successfully limits predator DNA

The method has been proven a feasible alternative and/or complement to visual inspection for analysis of fish stomach contents. It provided a comprehensive and detailed view of fish prey while successfully limiting the amplification and detection of predator DNA for all targeted species—the only exception was European sardine, from which a significant proportion of predator reads was obtained in real stomachs, although still lower than the proportion obtained in other studies not using BP and extracting the contents from the stomachs (Bessey et al., 2019; Piñol et al., 2014). Overall, the results obtained agree with previous knowledge: preference for arthropods by small pelagic fish (Bachiller et al., 2020; Bachiller & Irigoien, 2015) and dominance of fish DNA in the stomachs of European hake (Gül et al., 2023; Philips, 2012). We also identified overlap in the diet of pelagic species (Bachiller & Irigoien, 2015), and variation of prey composition in the stomachs of pelagic fish along the study area reflecting dietary plasticity and opportunistic feeding behaviour (Bachiller et al., 2020; Bogstad et al., 1995).

Yet, co-blocking of non-target species was observed for all the BP designed. Co-blocking occurs when amplification of other species than the target is prevented, and results in an artefactual decrease in the number of reads attributed to the co-blocked species (Piñol et al., 2015), ultimately leading to an underestimation of their contribution to the diet. However, co-blocking was typically limited, and the co-blocked species were still detected in the majority of mock samples even when using the lowest AP:BP ratios. Abstaining from using blocking primers to avoid co-blocking-derived bias will unarguably require manually extracting the stomach contents to maximise excluding predator DNA. Yet by doing so, the proportion of predator reads can easily reach 70-80% (Bessey et al., 2019; Piñol et al., 2014), resulting in less prey diversity detection and increased sample processing time.

### 4.2. Can DNA metabarcoding-based data contribute to trophic data collection?

Fish trophic data has been historically obtained through the visual inspection of stomach contents (e.g., López-López et al. (2012)). Accordingly, visually obtained data represents the main source of information on which ecosystem and multispecies models are based, and most models require input data to match visually derived ones. Yet, the emergence of methods other than visual inspection, such as stable isotopes or DNA metabarcoding, is providing additional information from which multispecies and ecosystem models could be greatly benefited (Hernvann et al., 2022; Matley et al., 2018). Below, we discuss how DNA metabarcoding-based data could potentially provide data to feed multispecies and ecosystems models by i) identifying fish trophic interactions, ii) quantifying the contribution of the different prey in the diet of predators, and iii) providing temporally and spatially comprehensive data.

#### 4.2.1. Identify trophic (prey-predator) interactions

DNA metabarcoding outperforms morphologically based approaches due to its higher detection and identification accuracy (e.g., Berry et al. (2015); Maes et al. (2022)). Consequently, it is expected to notably expand our knowledge on fish prey-predator interactions and thus provide a more accurate framework for ecosystems and multispecies models and for EBFM approaches. In particular, DNA metabarcoding is especially powerful for detecting and identifying usually overlooked prey by visual means, such as prey in early development stages, small plankton, or gelatinous organisms (Bachiller et al., 2020; Bachiller et al., 2021; Günther et al., 2021). Among these, gelatinous organisms are probably the most evident example of how DNA metabarcoding-based data can enhance understanding of the trophic web; based on DNA, over 20 cnidarian taxa were identified here as prey of the Atlantic mackerel, while none was reported by visual analysis. DNA metabarcoding is also a powerful tool to detect intraguild predation (Cuende et al., 2017), which is essential to determine fish natural mortality due to predation of early life stages that might be feeding on the same food resource as adults (Bachiller et al., 2015). Although based on genetic analyses we cannot determine whether the evidence of predation on a certain species corresponded to early life stages (more common and more feasible due to most predators’ size ranges) or adults (feasible for the largest fish, e.g., adult Atlantic mackerel or Atlantic horse mackerel), both options contribute to natural mortality of potential food competitors (Bachiller et al., 2015). We detected European anchovy and Atlantic horse mackerel in the stomach contents of the other pelagic species, European sardine in one Atlantic mackerel stomach, and Atlantic mackerel in one Atlantic horse mackerel stomach.

Yet, it should be acknowledged that DNA metabarcoding has also shortcomings; for instance, it is not able to identify cannibalism, a feeding strategy reported in fish (Bachiller et al., 2015; Pereira et al., 2017), and it cannot distinguish between primary and secondary consumption, i.e., DNA belonging to prey of prey (Traugott et al., 2021)—yet the molecular signal of secondary consumed prey is expected to be minimal since its DNA will represent a smaller proportion than directly ingested prey, and to be in a more advanced degradation state (Jakubavičiūtė et al., 2017). Complementary visual inspection of the stomach contents would be still needed to confirm cannibalism and/or whether a prey is more likely to come from direct or indirect consumption. Finally, is should be noted that the accuracy of DNA metabarcoding is highly dependent on the completeness of reference databases (Claver et al., 2023). In this sense, however, metabarcoding will only become a more powerful tool, as reference databases are being continuously populated and increased.

#### 4.2.2. Prey contribution in the diet of predators

Many multispecies and ecosystem models require prey contribution to be expressed as a proportion. DNA metabarcoding-derived data is thus *a priori* suitable input data for the models since it is expressed in proportion terms. However, read proportions in DNA metabarcoding are subjected to primer efficiency bias, which makes some groups or species to be more amplified than others so that read proportions may not represent actual prey proportions (Deagle et al., 2019; Piñol et al., 2015). Here, primer efficiency bias became evident in the mock assay—all species equally contributed to the mock samples in terms of DNA concentration but displayed divergent read proportions after the sequencing. However, this bias was not evident when comparing the diet composition of Atlantic mackerel assessed by both molecular and visual approaches, at least at higher taxonomic levels: both methods agreed pointing at Arthropoda and Vertebrata (fish) as primary food sources. The only exception was cnidarians, which represented up to 12.3% of Atlantic mackerel diet by DNA metabarcoding but were not detected visually. At this regard, it should be reminded that visual diet assessment is also subject to bias (Baker et al., 2014; Buckland et al., 2017; Moriarty et al., 2016), due to which highly digested contents, organisms that are rapidly degradable (e.g., gelatinous organisms), and some groups or early life stages especially difficult to identify visually are usually classified as unidentified prey or are not accounted. The usage of novel tools, such DNA metabarcoding and stable isotopes, is revealing widespread consumption of gelatinous plankton by fish (Bachiller et al., 2021; Günther et al., 2021), which suggests that the role of gelatinous organisms in marine trophic webs and in sustaining commercial fish populations may have been underestimated (Hays et al., 2018) and therefore overlooked by current food web and ecosystems models.

Although primer efficiency bias is a clear drawback for the straightforward incorporation of DNA metabarcoding-based data into ecosystem and multispecies models, it is noteworthy to remark that recent studies have proven that amplification efficiency can be measured based on mock assays, allowing the correction of read abundance values based on the observed primer efficiency bias making DNA metabarcoding a more reliable quantitative approach (Shelton et al., 2023; Silverman et al., 2021). Yet, how to construct mock samples assays for primer bias evaluation for the whole prey spectrum of fish in a given marine area is not an easy task.

#### 4.2.3. Temporally and spatially comprehensive data

Fish feeding behaviour is known to change temporally and spatially (Gül et al., 2023; Jansen et al., 2019; Olaso et al., 2005), and these variations should be accounted for by multispecies and ecosystems models to optimise the application of EBFM approaches—especially in the current context of climate change and poleward migration of many planktonic and nektonic species (Richardson, 2008), which is expected to impact currently established trophic webs. Our results indicate that DNA metabarcoding is capable of reporting such spatial and temporal dietary changes in great detail. This is exemplified here by variations in the diet composition of pelagic fish along the Bay of Biscay: the stomach contents of specimens captured by the same haul (i.e., in the same station) displayed more similar diet composition than specimens from different hauls, which is most likely reflecting prey availability. These results are in line with previous studies reporting opportunistic feeding behaviour and remarkable niche overlap in pelagic fish (Bachiller & Irigoien, 2015; Bogstad et al., 1995). Additionally, obtaining data on fish diet variations through time and space requires extensive, large-scaled stomach sampling, for which DNA metabarcoding represents a less time-consuming and more cost-effective alternative to traditional methods. Studies using DNA metabarcoding to detect temporal and spatial changes in fish diet, such as the one by Waraniak et al. (2019), are expected to provide a broader, more complete view of the dynamics of fish trophic interactions, which may ultimately facilitate the implementation of EBFM.

## 5. Concluding remarks

Implementing EBFM approaches requires extensive knowledge on trophic interactions, which is now more accessible thanks to emerging, alternative, or complementary methods for stomach content analysis. Here, we present a DNA metabarcoding-based method devised for speeding up the obtention of fish trophic data, suitable to be incorporated in routinary sampling programs thanks to the limited sample processing it requires. We show that the method is able to provide comprehensive lists of prey, accurate prey identification, and to report diet variations between predators reflecting species ecology and/or prey availability. In addition, thanks to the more cost-effectiveness of molecular methods compared to traditional analysis of stomach content, it is expected to decrease uncertainty in the models by allowing a notable increase of the sample size (Han et al., 2020). DNA metabarcoding still faces some drawbacks, due to which the usage of this kind of data by the models is not straightforward. Some of them can be overcome, such as improving the confidence on the quantitative value of DNA metabarcoding relative read abundances (Shelton et al., 2023; Silverman et al., 2021); others will remain unsolvable, such as identifying the stage development, age, or length of the prey, detecting cannibalism, or distinguishing between primary and secondary consumption, for which visual inspection would still be needed. In any case, regardless of its limitations and biases, DNA metabarcoding-derived data is expected to be useful for ecosystem and multispecies models, especially considering that visual diet assessment is also subject to bias (Baker et al., 2014; Buckland et al., 2017; Moriarty et al., 2016). Instead of disregarding any method, identify and combine the strengths of each method is probably the best approach for feeding multispecies and ecosystem models and, ultimately, for a broader and more efficient implementation of EBFM (Hernvann et al., 2022; Matley et al., 2018).

## Funding Information

This study was funded by the projects GENGES (through a framework contract with the Department of Agriculture and Fisheries of the Basque Government) and TROGEN (through the EMFF (European Maritime Fishing Funds) Data collection framework). XC was supported by the Spanish National Program Juan de la Cierva-Formación (MICINN / AEI / 726 10.13039/501100011033 FJC2020-044367-I) funded by the Government of Spain (Ministry of Science and Innovation (MICINN) and the State Research Agency (AEI) and the European Union (NextGenerationEU/PRTR).

## Acknowledgements

We would like to thank Paula Alvárez, Maria Santos, Guillermo Boyra, Iñaki Quincoces, and Jesus Martinez for their help in obtaining stomach samples during during the TRIENAL, BIOMAN, JUVENA and ITSASTEKA surveys scientific surveys. This study was funded by the projects GENGES (through a framework contract with the Department of Agriculture and Fisheries of the Basque Government) and TROGEN (through the EMFF (European Maritime Fishing Funds) Data collection framework). XC was supported by the Spanish National Program Juan de la Cierva-Formación (MICINN / AEI / 726 10.13039/501100011033 FJC2020-044367-I) funded by the Government of Spain (Ministry of Science and Innovation (MICINN) and the State Research Agency (AEI) and the European Union (NextGenerationEU/PRTR).

## Conflict of interest

Authors declare no conflict of interest.

## Authors contributions

UC, IM and NRE designed research. OC, AL, IM, EB, and NRE performed research. OC, AL, IM, UC, and NRE contributed new reagents or analytical tools. OC wrote the paper, with insightful contributions from EA, XC, AL, UC, EB and NRE. All authors revised the manuscript and agreed with its publication.

## Notes

### Competing Interest Statement

The authors have declared no competing interest.

