## Supplementary material for "Increasing marine trophic web knowledge through DNA analyses of fish stomach contents: a step towards an Ecosystem Based Approach to fisheries research": Figure S1

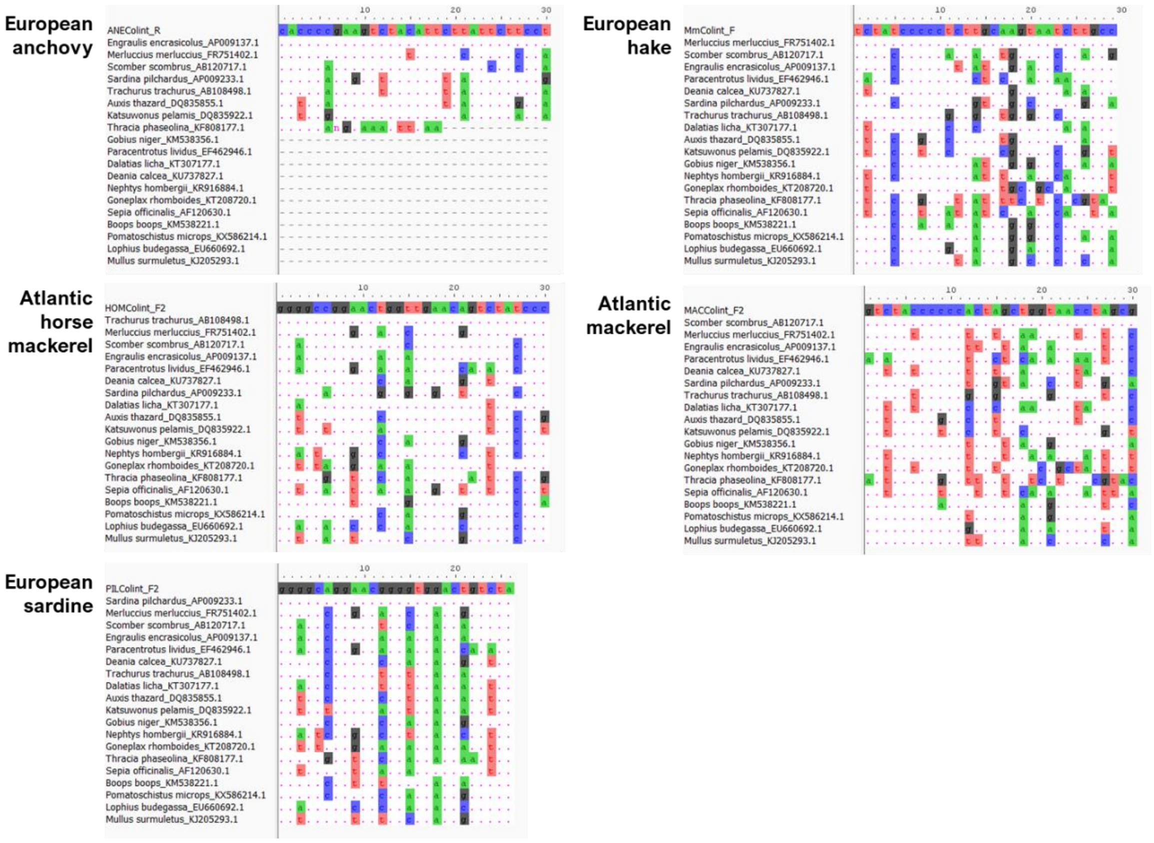


**Figure S1.** Alignment showing the mismatches between the BPs for European anchovy (A*), Atlantic horse mackerel (B), European sardine (C), European hake (D), and Atlantic mackerel (E), and the sequences of the species included in the mock samples (GenBank accession number of the sequences used are specified just after the species name). * For European anchovy, most of the references available in GenBank were not long enough to cover the BP region.
