## Supplementary material for "Increasing marine trophic web knowledge through DNA analyses of fish stomach contents: a step towards an Ecosystem Based Approach to fisheries research": Table S1

**Table S1.** List of the 21 species conforming the mock stomachs generated for testing the efficiency of the blocking primers. * indicate the predator species targeted in the study.

| Species | Taxonomy (Class, Phylum) |
|---|---|
| Engraulis encrasicolus* | Actinopterygii, Chordata |
| Merluccius merluccius* | Actinopterygii, Chordata |
| Sardina pilchardus* | Actinopterygii, Chordata |
| Scomber scombrus* | Actinopterygii, Chordata |
| Trachurus trachurus* | Actinopterygii, Chordata |
| Boops boops | Actinopterygii, Chordata |
| Lophius budegassa | Actinopterygii, Chordata |
| Mullus surmuletus | Actinopterygii, Chordata |
| Auxis thazard | Actinopterygii, Chordata |
| Katsuwonus pelamis | Actinopterygii, Chordata |
| Gobius niger | Actinopterygii, Chordata |
| Pomatoschistus microps | Actinopterygii, Chordata |
| Deania calcea | Elasmobranchii, Chordata |
| Scymnodon ringens | Elasmobranchii, Chordata |
| Dalatias licha | Elasmobranchii, Chordata |
| Nephtys hombergii | Annelida, Polychaeta |
| Glycera lapidum | Annelida, Polychaeta |
| Goneplax rhomboides | Crustacea, Arthropoda |
| Paracentrotus lividus | Echinoidea, Echinodermata |
| Thracia phaseolina | Bivalbia, Mollusca |
| Sepia officinalis | Cephalopoda, Mollusca |
