## Supplementary material for "Increasing marine trophic web knowledge through DNA analyses of fish stomach contents: a step towards an Ecosystem Based Approach to fisheries research": Table S2

**Table S2:** Detailed information on the specimens analysed in the study, including the oceanographic survey during which the specimens were captured, the date and coordinates of capture, the length and weight of the individuals, the haul during which the specmines were obtained, and the AP:BP ratio used for the stomach contents amplification. Latitude and longitude coordinates refers to the initial point of the haul transect.

| Species | Sample_ID | Survey | Capture date | Lat | Long | Haul | Length (mm) | Weight (g) | AP:BP ratio |
|---|---|---|---|---|---|---|---|---|---|
| European hake (*Merluccius merluccius*) | HKE_01 | Itsasteka 2013 | 25/06/2013 | 43.53 | -2.47 | ITS_1 | 210 | 69 | 1:5 |
| | HKE_02 | Itsasteka 2013 | 25/06/2013 | 43.53 | -2.47 | ITS_1 | 185 | 37 | 1:5 |
| | HKE_03 | Itsasteka 2013 | 25/06/2013 | 43.53 | -2.47 | ITS_1 | 200 | 44 | 1:5 |
| | HKE_04 | Itsasteka 2013 | 25/06/2013 | 43.53 | -2.47 | ITS_1 | 210 | 63 | 1:5 |
| | HKE_05 | Itsasteka 2013 | 25/06/2013 | 43.53 | -2.47 | ITS_1 | 230 | 74 | 1:5 |
| | HKE_06 | Itsasteka 2013 | 25/06/2013 | 43.53 | -2.47 | ITS_1 | 425 | 555 | 1:5 |
| | HKE_07 | Itsasteka 2013 | 25/06/2013 | 43.53 | -2.47 | ITS_1 | 205 | 54 | 1:5 |
| | HKE_08 | Itsasteka 2013 | 25/06/2013 | 43.53 | -2.47 | ITS_1 | 405 | 475 | 1:5 |
| European anchovy (*Engraulis encrasicolus*) | ANE_11 | JUVENA 2014 | 11/09/2014 | 43.86 | -1.77 | JUV_9012 | 125 | 14.6 | 1:2.5 |
| | ANE_12 | JUVENA 2014 | 11/09/2014 | 43.86 | -1.77 | JUV_9012 | 125 | 13.5 | 1:2.5 |
| | ANE_14 | JUVENA 2014 | 11/09/2014 | 43.86 | -1.77 | JUV_9012 | 125 | 14.3 | 1:2.5 |
| | ANE_15 | JUVENA 2014 | 11/09/2014 | 43.86 | -1.77 | JUV_9012 | 125 | 15.1 | 1:2.5 |
| | ANE_17 | JUVENA 2014 | 11/09/2014 | 43.86 | -1.77 | JUV_9012 | 125 | 14.9 | 1:2.5 |
| | ANE_22 | JUVENA 2014 | 19/09/2014 | 47.38 | -3.64 | JUV_9025 | 154 | 23.7 | 1:2.5 |
| European sardine (*Sardina pilchardus*) | PIL_01 | JUVENA 2014 | 11/09/2014 | 43.86 | -1.77 | JUV_9012 | 193 | 58.2 | 1:2.5 |
| | PIL_07 | JUVENA 2014 | 11/09/2014 | 43.86 | -1.77 | JUV_9012 | 190 | 52.3 | 1:2.5 |
| | PIL_08 | JUVENA 2014 | 11/09/2014 | 43.86 | -1.77 | JUV_9012 | 186 | 52.8 | 1:2.5 |
| | PIL_09 | JUVENA 2014 | 11/09/2014 | 43.86 | -1.77 | JUV_9012 | 186 | 47.5 | 1:2.5 |
| | PIL_10 | JUVENA 2014 | 11/09/2014 | 43.86 | -1.77 | JUV_9012 | 190 | 42.3 | 1:2.5 |
| | PIL_11 | JUVENA 2014 | 19/09/2014 | 47.50 | -3.37 | JUV_9026 | 161 | 36.1 | 1:2.5 |
| | PIL_13 | JUVENA 2014 | 19/09/2014 | 47.50 | -3.37 | JUV_9026 | 168 | 38.4 | 1:2.5 |
| | PIL_17 | JUVENA 2014 | 19/09/2014 | 47.50 | -3.37 | JUV_9026 | 178 | 45 | 1:2.5 |
| | PIL_20 | JUVENA 2014 | 19/09/2014 | 47.50 | -3.37 | JUV_9026 | 234 | 83.4 | 1:2.5 |
| Atlantic horse mackerel (*Trachurus trachurus*) | HOM_08 | JUVENA 2014 | 11/09/2014 | 43.86 | -1.77 | JUV_9012 | 143 | 23.1 | 1:10 |
| | HOM_11 | JUVENA 2014 | 19/09/2014 | 47.50 | -3.37 | JUV_9026 | 98.00 | 8.00 | 1:10 |
| | HOM_11_R | JUVENA 2014 | 19/09/2014 | 47.50 | -3.37 | JUV_9026 | 98.00 | 8.00 | 1:5 |
| | HOM_13 | JUVENA 2014 | 19/09/2014 | 47.50 | -3.37 | JUV_9026 | 111.00 | 11.40 | 1:10 |
| | HOM_13_R | JUVENA 2014 | 19/09/2014 | 47.50 | -3.37 | JUV_9026 | 111.00 | 11.40 | 1:5 |
| | HOM_16 | JUVENA 2014 | 19/09/2014 | 47.50 | -3.37 | JUV_9026 | 158.00 | 31.10 | 1:10 |
| | HOM_16_R | JUVENA 2014 | 19/09/2014 | 47.50 | -3.37 | JUV_9026 | 158.00 | 31.10 | 1:5 |
| | HOM_17 | JUVENA 2014 | 19/09/2014 | 47.50 | -3.37 | JUV_9026 | 152.00 | 28.30 | 1:10 |
| | HOM_17_R | JUVENA 2014 | 19/09/2014 | 47.50 | -3.37 | JUV_9026 | 152.00 | 28.30 | 1:5 |
| | HOM_23 | JUVENA 2014 | 27/09/2014 | 43.78 | -7.17 | JUV_9032 | 232.00 | 108.80 | 1:10 |
| | HOM_23_R | JUVENA 2014 | 27/09/2014 | 43.78 | -7.17 | JUV_9032 | 232.00 | 108.80 | 1:5 |
| Atlantic mackerel (*Scomber scombrus*) - adults | MAC_01 | JUVENA 2014 | 11/09/2014 | 43.86 | -1.77 | JUV_9012 | 185.00 | 57.10 | 1:10 |
| | MAC_01_R | JUVENA 2014 | 11/09/2014 | 43.86 | -1.77 | JUV_9012 | 185.00 | 57.10 | 1:5 |
| | MAC_03 | JUVENA 2014 | 11/09/2014 | 43.86 | -1.77 | JUV_9012 | 167.00 | 37.50 | 1:10 |
| | MAC_03_R | JUVENA 2014 | 11/09/2014 | 43.86 | -1.77 | JUV_9012 | 167.00 | 37.50 | 1:5 |
| | MAC_05 | JUVENA 2014 | 11/09/2014 | 43.86 | -1.77 | JUV_9012 | 170 | 44.6 | 1:10 |
| | MAC_07 | JUVENA 2014 | 11/09/2014 | 43.86 | -1.77 | JUV_9012 | 171 | 44.7 | 1:10 |
| | MAC_09 | JUVENA 2014 | 11/09/2014 | 43.86 | -1.77 | JUV_9012 | 166 | 37.1 | 1:10 |
| | MAC_10 | JUVENA 2014 | 11/09/2014 | 43.86 | -1.77 | JUV_9012 | 163.00 | 34.30 | 1:10 |
| | MAC_10_R | JUVENA 2014 | 11/09/2014 | 43.86 | -1.77 | JUV_9012 | 163.00 | 34.30 | 1:5 |
| | MAC_12 | JUVENA 2014 | 19/09/2014 | 47.50 | -3.37 | JUV_9026 | 193 | 50.5 | 1:10 |
| | MAC_14 | JUVENA 2014 | 19/09/2014 | 47.50 | -3.37 | JUV_9026 | 195 | 50.5 | 1:10 |
| | MAC_15 | JUVENA 2014 | 19/09/2014 | 47.50 | -3.37 | JUV_9026 | 199 | 59 | 1:10 |
| | MAC_18 | JUVENA 2014 | 19/09/2014 | 47.50 | -3.37 | JUV_9026 | 195.00 | 48.60 | 1:10 |
| | MAC_18_R | JUVENA 2014 | 19/09/2014 | 47.50 | -3.37 | JUV_9026 | 195.00 | 48.60 | 1:5 |
| | MAC_23 | JUVENA 2014 | 27/09/2014 | 43.78 | -7.17 | JUV_9032 | 200 | 60.9 | 1:10 |
| | MAC_27 | JUVENA 2014 | 27/09/2014 | 43.78 | -7.17 | JUV_9032 | 202 | 62.5 | 1:10 |
| | MAC_28 | JUVENA 2014 | 27/09/2014 | 43.78 | -7.17 | JUV_9032 | 205 | 65.3 | 1:10 |
| | MAC_29 | JUVENA 2014 | 27/09/2014 | 43.78 | -7.17 | JUV_9032 | 202 | 60.1 | 1:10 |

| | | | | | | | | | |
|---|---|---|---|---|---|---|---|---|---|
| Atlantic mackerel (*Scomber scombrus* ) adults | 16MAC_01 | TRIENAL 2016 | 20/03/2016 | 46.23 | -3.07 | TRI_1 | 293 | 146.9 | 1:10 |
| | 16MAC_02 | TRIENAL 2016 | 20/03/2016 | 46.23 | -3.07 | TRI_1 | 330 | 218.2 | 1:10 |
| | 16MAC_03 | TRIENAL 2016 | 20/03/2016 | 46.23 | -3.07 | TRI_1 | 360 | 308.8 | 1:10 |
| | 16MAC_04 | TRIENAL 2016 | 20/03/2016 | 46.23 | -3.07 | TRI_1 | 355 | 278.2 | 1:10 |
| | 16MAC_05 | TRIENAL 2016 | 20/03/2016 | 46.23 | -3.07 | TRI_1 | 340 | 274.2 | 1:10 |
| | 16MAC_07 | TRIENAL 2016 | 20/03/2016 | 46.23 | -3.07 | TRI_1 | 347 | 273.8 | 1:10 |
| | 16MAC_08 | TRIENAL 2016 | 20/03/2016 | 46.23 | -3.07 | TRI_1 | 344 | 256.5 | 1:10 |
| | 16MAC_09 | TRIENAL 2016 | 20/03/2016 | 46.23 | -3.07 | TRI_1 | 395 | 343.4 | 1:10 |
| | 16MAC_10 | TRIENAL 2016 | 20/03/2016 | 46.23 | -3.07 | TRI_1 | 294 | 164.3 | 1:10 |
| | 16MAC_14 | TRIENAL 2016 | 25/03/2016 | 46.05 | -4.66 | TRI_3 | 347 | 257.2 | 1:10 |
| | 16MAC_16 | TRIENAL 2016 | 25/03/2016 | 46.05 | -4.66 | TRI_3 | 382 | 363.8 | 1:10 |
| | 16MAC_17 | TRIENAL 2016 | 25/03/2016 | 46.05 | -4.66 | TRI_3 | 336 | 238.1 | 1:5 |
| | 16MAC_18 | TRIENAL 2016 | 25/03/2016 | 46.05 | -4.66 | TRI_3 | 366 | 304.8 | 1:5 |
| | 16MAC_19 | TRIENAL 2016 | 25/03/2016 | 46.05 | -4.66 | TRI_3 | 361 | 293.5 | 1:10 |
| | 16MAC_20 | TRIENAL 2016 | 25/03/2016 | 46.05 | -4.66 | TRI_3 | 345 | 292.4 | 1:10 |
| | 16MAC_21 | TRIENAL 2016 | 05/04/2016 | 45.25 | -3.42 | TRI_5 | 372 | 334 | 1:5 |
| | 16MAC_22 | TRIENAL 2016 | 05/04/2016 | 45.25 | -3.42 | TRI_5 | 328 | 232.4 | 1:5 |
| | 16MAC_24 | TRIENAL 2016 | 05/04/2016 | 45.25 | -3.42 | TRI_5 | 324 | 207 | 1:5 |
| | 16MAC_31 | TRIENAL 2016 | 05/04/2016 | 45.75 | -3.39 | TRI_6 | 340 | 249 | 1:5 |
| | 16MAC_34 | TRIENAL 2016 | 05/04/2016 | 45.75 | -3.39 | TRI_6 | 318 | 219.9 | 1:10 |
| | 16MAC_35 | TRIENAL 2016 | 05/04/2016 | 45.75 | -3.39 | TRI_6 | 340 | 277.9 | 1:10 |
| | 16MAC_36 | TRIENAL 2016 | 05/04/2016 | 45.75 | -3.39 | TRI_6 | 336 | 253.2 | 1:10 |
| | 16MAC_37 | TRIENAL 2016 | 05/04/2016 | 45.75 | -3.39 | TRI_6 | 374 | 316 | 1:10 |
| | 16MAC_38 | TRIENAL 2016 | 05/04/2016 | 45.75 | -3.39 | TRI_6 | 352 | 269.2 | 1:3 |
| | 16MAC_39 | TRIENAL 2016 | 05/04/2016 | 45.75 | -3.39 | TRI_6 | 350 | 266.6 | 1:5 |
| | 17MAC01 | BIOMAN 2016 | 15/05/2016 | 44.10 | -1.71 | BIO_5017 | 375 | 339.9 | 1:5 |
| | 17MAC02 | BIOMAN 2016 | 15/05/2016 | 44.10 | -1.71 | BIO_5017 | 380 | 416.78 | 1:5 |
| | 17MAC04 | BIOMAN 2016 | 15/05/2016 | 44.10 | -1.71 | BIO_5017 | 393 | 397.28 | 1:5 |
| | 17MAC05 | BIOMAN 2016 | 15/05/2016 | 44.10 | -1.71 | BIO_5017 | 363 | 374.1 | 1:5 |
| | 17MAC06 | BIOMAN 2016 | 15/05/2016 | 44.10 | -1.71 | BIO_5017 | 381 | 396.38 | 1:5 |
| | 17MAC07 | BIOMAN 2016 | 15/05/2016 | 44.10 | -1.71 | BIO_5017 | 343 | 309.62 | 1:5 |
| | 17MAC08 | BIOMAN 2016 | 15/05/2016 | 44.38 | -1.97 | BIO_5018 | 396 | 442.57 | 1:3 |
| | 17MAC09 | BIOMAN 2016 | 15/05/2016 | 44.38 | -1.97 | BIO_5018 | 371 | 363.79 | 1:5 |
| | 17MAC11 | BIOMAN 2016 | 15/05/2016 | 44.38 | -1.97 | BIO_5018 | 367 | 322.09 | 1:3 |
| | 17MAC12 | BIOMAN 2016 | 15/05/2016 | 44.38 | -1.97 | BIO_5018 | 361 | 289.94 | 1:5 |
| | 17MAC16 | BIOMAN 2016 | 15/05/2016 | 44.38 | -1.97 | BIO_5018 | 372 | 361.25 | 1:5 |
| | 17MAC18 | BIOMAN 2016 | 20/05/2016 | 44.45 | -1.41 | BIO_5027 | 251 | 110.71 | 1:5 |
| | 17MAC20 | BIOMAN 2016 | 20/05/2016 | 44.45 | -1.41 | BIO_5027 | 402 | 412.22 | 1:5 |
| | 17MAC30 | BIOMAN 2016 | 20/05/2016 | 44.74 | -1.41 | BIO_5028 | 263 | 135.83 | 1:3 |
| | 17MAC42 | BIOMAN 2016 | 25/05/2016 | 46.41 | -2.51 | BIO_5038 | 327 | 249.97 | 1:5 |
| | 17MAC45 | BIOMAN 2016 | 25/05/2016 | 46.41 | -2.51 | BIO_5038 | 338 | 261.68 | 1:5 |
| Atlantic mackerel juveniles | 17MAC49 | BIOMAN 2016 | 17/05/2016 | 46.00 | -2.79 | BIO_MIK36 | 67 | 1.11 | 1:5 |
| | 17MAC50 | BIOMAN 2016 | 17/05/2016 | 46.00 | -2.79 | BIO_MIK36 | 74 | 1.479 | 1:5 |
| | 17MAC51 | BIOMAN 2016 | 17/05/2016 | 46.00 | -2.79 | BIO_MIK36 | 63 | 0.779 | 1:5 |
| Atlantic mackerel larvae | LAR_01_16 | TRIENAL 2013 | 25/03/2013 | 46.75 | -4.91 | N/A | N/A | N/A | 1:2 |
| | LAR_01_20 | TRIENAL 2013 | 25/03/2013 | 46.75 | -4.91 | N/A | N/A | N/A | 1:2 |
| | LAR_01_21 | TRIENAL 2013 | 25/03/2013 | 46.75 | -4.91 | N/A | N/A | N/A | 1:2 |
| | LAR_ST1_13 | Careva0313 | 14/03/2013 | 43.43 | -3.15 | N/A | N/A | N/A | 1:2 |
| | LAR_ST2_02 | Careva0313 | 15/03/2013 | 43.63 | -3.83 | N/A | N/A | N/A | 1:2 |
