## Supplementary material for "Increasing marine trophic web knowledge through DNA analyses of fish stomach contents: a step towards an Ecosystem Based Approach to fisheries research": Table S3

**Table S3.** Experimental design for assessing the efficiency of the generated BPs in mock stomachs. For each species, different AP:BP ratios and different prey DNA proportion (prey %) were used.

| AP:BP ratio | prey % | Replicate |
|---|---|---|
| no BP | 100 | 1 |
| no BP | 100 | 2 |
| 1:2 | 100 | 1 |
| 1:2 | 100 | 2 |
| 1:5 | 100 | 1 |
| 1:5 | 100 | 2 |
| 1:10 | 100 | 1 |
| 1:10 | 100 | 2 |
| no BP | 70 | 1 |
| no BP | 70 | 2 |
| 1:2 | 70 | 1 |
| 1:2 | 70 | 2 |
| 1:5 | 70 | 1 |
| 1:5 | 70 | 2 |
| 1:10 | 70 | 1 |
| 1:10 | 70 | 2 |
| no BP | 30 | 1 |
| no BP | 30 | 2 |
| 1:2 | 30 | 1 |
| 1:2 | 30 | 2 |
| 1:5 | 30 | 1 |
| 1:5 | 30 | 2 |
| 1:10 | 30 | 1 |
| 1:10 | 30 | 2 |
| no BP | 0 | 1 |
| no BP | 0 | 2 |
| 1:2 | 0 | 1 |
| 1:2 | 0 | 2 |
| 1:5 | 0 | 1 |
| 1:5 | 0 | 2 |
| 1:10 | 0 | 1 |
| 1:10 | 0 | 2 |
